# Majority of human circulating plasmablasts stop blasting: A probable misnomer

**DOI:** 10.1101/2023.09.10.557057

**Authors:** Doan C. Nguyen, Celia Saney, Ian T. Hentenaar, Monica Cabrera-Mora, Matthew C. Woodruff, Joel Andrews, Sagar Lonial, Ignacio Sanz, F. Eun-Hyung Lee

**Affiliations:** Division of Pulmonary, Allergy, Critical Care, and Sleep Medicine, Department of Medicine, Emory University, Atlanta, GA, United States; Division of Rheumatology, Department of Medicine, Emory University, Atlanta, GA, United States; Lowance Center for Human Immunology, Emory University, Atlanta, GA, United States; Department of Hematology and Medical Oncology, Winship Cancer Institute, Emory University, Atlanta, GA, United States

## Abstract

Following infection or vaccination, early-minted antibody secreting cells (ASC) or plasmablasts appear in circulation transiently, and a small fraction migrates to the spleen or bone marrow (BM) to mature into long-lived plasma cells (LLPC). While LLPC, by definition, are quiescent or non-dividing, the majority of blood ASC are thought to be “blasting” or proliferative. In this study, we find >95% nascent blood ASC in culture express Ki-67 but only 6-12% incorporate BrdU after 4h or 24h labeling. In contrast, <5% BM LLPC in culture are Ki-67^+^ with no BrdU uptake. Due to limitations of traditional flow cytometry, we utilized a novel optofluidic technology to evaluate cell division with simultaneous functional Ig secretion. We find 11% early-minted blood ASC undergo division, and none of the terminally differentiated BM LLPC (CD19^-^CD38^hi^CD138^+^) divide during the 7-21 days in culture. While BM LLPC undergo complete cell cycle arrest, the process of differentiation into an ASC of plasmablasts discourages entry into S phase. Since the majority of Ki-67^+^ nascent blood ASC have exited cell cycle and are no longer actively “blasting”, the term “plasmablast”, which traditionally refers to an ASC that still has the capacity to divide, may probably be a misnomer.

## Introduction

During infection or vaccination, B cells undergo proliferation and differentiation to generate protective antibody responses. Activation of the B cell receptors (BCR) following antigen encounter triggers a cascade of signaling events that lead to proliferation of B cells^1^. In addition to BCR and toll-like receptors (TLR), cytokine receptors and TNF family receptors, such as CD40^2,3^, also enhance B cell proliferation. Upon induction, the first division of a naїve B cell requires 24 hours (h), and the cells continue to divide every 6-8h thereafter prior to differentiating into antibody secreting cells (ASC) or plasmablasts^4,5^. The term “plasmablast” has traditionally been used interchangeably with early-minted ASC with proliferative potential and ongoing cell division^6-11^. While the majority of ASC undergo apoptosis, a small fraction migrates to bone marrow (BM) and further matures into long-lived plasma cells (LLPC)^12,13^. BM LLPC have extraordinary longevity and provide durable humoral protection against reinfection for a lifetime^14,15^. By definition, BM LLPC are non-dividing, terminally differentiated cells^16^.

Commonly used methods for measuring cell proliferation include expression of Ki-67, a nuclear protein strongly expressed in proliferating cells (although its function is unclear), and incorporation of thymidine analogs (BrdU) into the DNA of actively dividing cells. Studies interrogating intracellular Ki-67 expression or BrdU incorporation have suggested human BM LLPC do not divide while ASC in the blood are proliferative^15,17,18^. In non-human primates, BrdU^+^ BM LLPC can be found a decade after BrdU administration^19^, suggesting persistence for years in the BM of the same ASC (i.e. without division). In contrast, the majority (>70-98%) of nascent blood ASC in vaccinated individuals are Ki-67^+15,17,20^. Whether these early-minted ASC have recently divided or if they are still actively cycling upon entering the bloodstream remains incompletely understood.

In this study, we overcome limitations in ASC survival *ex vivo* with a new human *in vitro* plasma cell survival system (PCSS)^21,22^. These culture systems allow us to visualize single ASC proliferation using a novel optofluidic platform to measure cell division simultaneously with ongoing Ig secretion. Results show that 90% of early-minted blood ASC stop dividing and that BM LLPC remain in a state of permanent cell cycle arrest. These results emphasize induction of networks that promote cell cycle arrest in early ASC differentiation.

## Results

### Few blood ASC incorporate BrdU or undergo proliferation

Human early-minted ASC (CD19^lo^CD27^hi^CD38^hi^) were isolated from the blood by FACS sorting from healthy adults between 5-7 days after vaccination (since days 5-7 post-vaccination reveal the peak window for antigen-specific ASC circulation^23-27^). As previously shown^15,17,20^, the majority of blood ASC are Ki-67^+^ even when cultured in PCSS: **Fig 1a,b** shows that blood ASC express Ki-67 as high as 99% (99.1±0.8%), 97% (97.3±1.3%), and 95% (95.1±1.7%) on day 0, 1, and 3, respectively. For non-proliferative control, we used unstimulated blood naive B cells. These cultured ASC also secrete IgG when assayed by bulk ELISpots (**Suppl. Fig. S1a,b**).

**Fig. 1.**
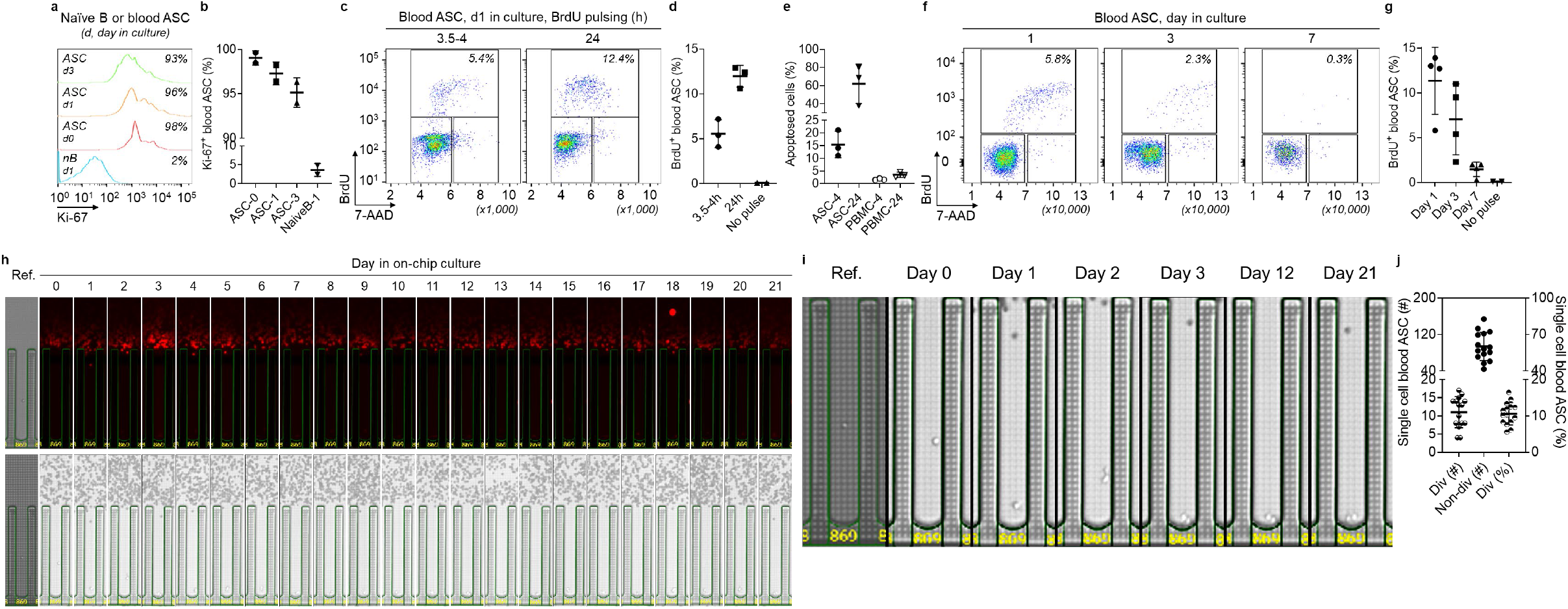
Very few early-minted ASC (plasmablasts) divide. (**a**) Representative flow cytometric analysis of Ki-67 expression in blood ASC at day 0, 1, and 3 in culture (with unstimulated naїve B cells at day 1 as a non-proliferative control). The numbers indicate the percentages of the cells positive for Ki-67. (**b**) Quantitation of Ki-67^+^ blood ASC (n=2 independent biological replicates). 0, 1, 3, day in culture. (**c**) Representative flow cytometric analysis of BrdU incorporation in blood ASC at day 1 in culture after 3.5-4h or 24h pulsing. (**d**) Quantitation of BrdU^+^ blood ASC (n=3 independent biological replicates). h, hours BrdU pulsed. All cells were assayed at day 1 in culture. (**e**) Quantitation of apoptosed blood ASC after 3.5-4h or 24h BrdU pulsing (n=3 independent biological replicates). PBMC were used as control populations. 4, 24, hours BrdU pulsed. All cells were assayed at day 1 in culture. (**f**) Representative flow cytometric analysis of BrdU incorporation in blood ASC at day 1, 3, and 7 in culture after 3.5-4h pulsing (with apoptotic cells excluded). (**g**) Quantitation of BrdU^+^ blood ASC (n=4 independent biological replicates). All 3.5-4h BrdU pulsed. In (**c,f**): Individual cell cycle phases (S, BrdU^+^; G0/G1, 2N and BrdU^-^; and G2/M, 4N and BrdU^-^) were indicated and the percentage of ASC in S phase was estimated by the amount of BrdU detected. Apoptotic cells were excluded from the analysis. (**h**) Image series displaying a single ASC with ongoing Ig secretion showing evidence of division (bright field) from day 0 to 21. (**i**) Zoomed in bright-field image series for direct visualization of the dividing cells from (**h**) on select days in culture. (**j**) Quantitation (left y axis) and frequency (right y axis) of dividing and non-dividing single cell blood ASC across 15 independent experiments. Div, dividing; Non-div, no dividing. In (**h,i**): Ref., reference image for visualization of the number of cells penned at loading.

To assess the phases of cell cycle, we employed flow cytometric analysis of BrdU incorporation in blood ASC (**Suppl. Fig. S1c**). After a 3.5-4h BrdU pulse label of blood ASC, we show 6% (5.6±1.6%) the cells were in S phase on day 1 of culture (**Fig. 1c**, left panel, and **Fig. 1d**). In an attempt to capture asynchronous cells beyond the 4h pulse, we labeled for 24h and found 12% (12±1.2%) of cells in S phase (**Fig. 1c**, right panel and **Fig. 1d**). Notably, the portion of cell death was substantially higher when pulsing for 24h (61%; 61.4±21.1%) compared to that of 4h (15%; 15.5±5.2%) (**Fig. 1e**). These results suggest longer BrdU labeling times may be toxic to blood ASC in culture with potential overestimation of labeled cells due to increased cell death. Ultimately, longer labeling may actually lead to less predictable cell recovery and viability.

As 4h pulses appeared to be sufficient to balance BrdU incorporation and toxicity for early-minted blood ASC, we applied it for all subsequent BrdU experiments. We found that the frequency of blood ASC in S phase decreased from 11% (11.4±3.7%) on day 1 to 7% (7.1±3.9%) on day 3, and further dropped to 1% (1.5±0.8%) by day 7 (**Fig. 1f,g**). By day 22, S phase entry was not detected in blood ASC despite functional IgG secretion (**Suppl. Fig. S1 d,e,f**), indicating that nascent ASC in long-term culture stop proliferating. These data suggest 90% early-minted ASC in the culture exit cell cycle early (at days 1-3) and nearly all are in complete cell cycle arrest from day 7 onwards.

Due to cell toxicity with longer pulses, BrdU labeling was not ideal to assess proliferation rates of ASC accurately. To overcome these issues, we evaluated ASC cell division coupled with ongoing IgG secretion by single cell analysis. We loaded early-minted blood ASC onto the optofluidic Lightning platform (Berkeley Lights, Inc.; now PhenomeX, Inc.) and measured IgG blooms and counted cell numbers by bright field each day for 7-21 days in PCSS cultures. We observed by day 15, the majority of ongoing IgG secreting cells (IgG-blooming pens) had not divided (**Suppl. Fig. S2a**). Only 11% (10.5±3.1%) had undergone cell division (**Fig. 1h-j**).

While some division could occur as early as 8h (**Suppl. Fig. S2b**) if cells divided, the division occurred within 24-48h (**Fig. 1h,i** and **Suppl. Fig. S2c**). Even when followed for up to 21 days, no further divisions were observed (**Fig. 1h**). Importantly, during these time periods, the lack of the first or any potential additional cell proliferation was not due to cell death, since intact cells were visualized with effective ongoing IgG blooms as late as day 21 (**Fig. 1h**). These results demonstrate that only 11% of *bona fide* IgG ASC in culture undergo proliferation over the observation course of up to 21 days.

### BM LLPC do not undergo cell division

In direct contrast to the blood ASC, human BM LLPC, defined as CD19^-^CD38^+^CD138^+^ ASC (PopD)^12,15^, retain long-lived antigen specificities from exposures over 40 years and have been shown to be non-dividing by Ki-67 expression^15,17,18^. Similar to previous studies, BM LLPC demonstrate only <5% Ki-67 staining compared to total BM mononuclear cells (BMMC) showing 44% (**Fig. 2a,b** and **Suppl. Fig. S3a**). For non-proliferative and proliferative control, we used unstimulated naive B cells and BMMC, respectively. We further confirmed that BM LLPC cultured in PCSS were indeed functionally secreting IgG (**Suppl. Fig. 3b,c**) despite minimal intracellular Ki-67 expression: 5.2% (5.2±2.26%) and 5.1% (5.1±1.03%) on day 0 and 1, respectively. To assess the phases of cell cycle, we employed flow cytometric analysis of BrdU incorporation in BM LLPC (**Suppl. Fig. S3d**). After BrdU pulse label for 4h on day 1, 3, and 7 in culture; we observed <0.1% BrdU incorporation at all time points (**Fig. 2c,d**). Finally, to assess if BM LLPC in culture could undergo division beyond day 7, we bulk-cultured BM LLPC for 56 days and found negligible Ki-67 expression or BrdU incorporation (**Suppl. Fig. S3e,f**), despite ongoing ability to secrete IgG (**Suppl. Fig. S3g,h**).

**Fig. 2.**
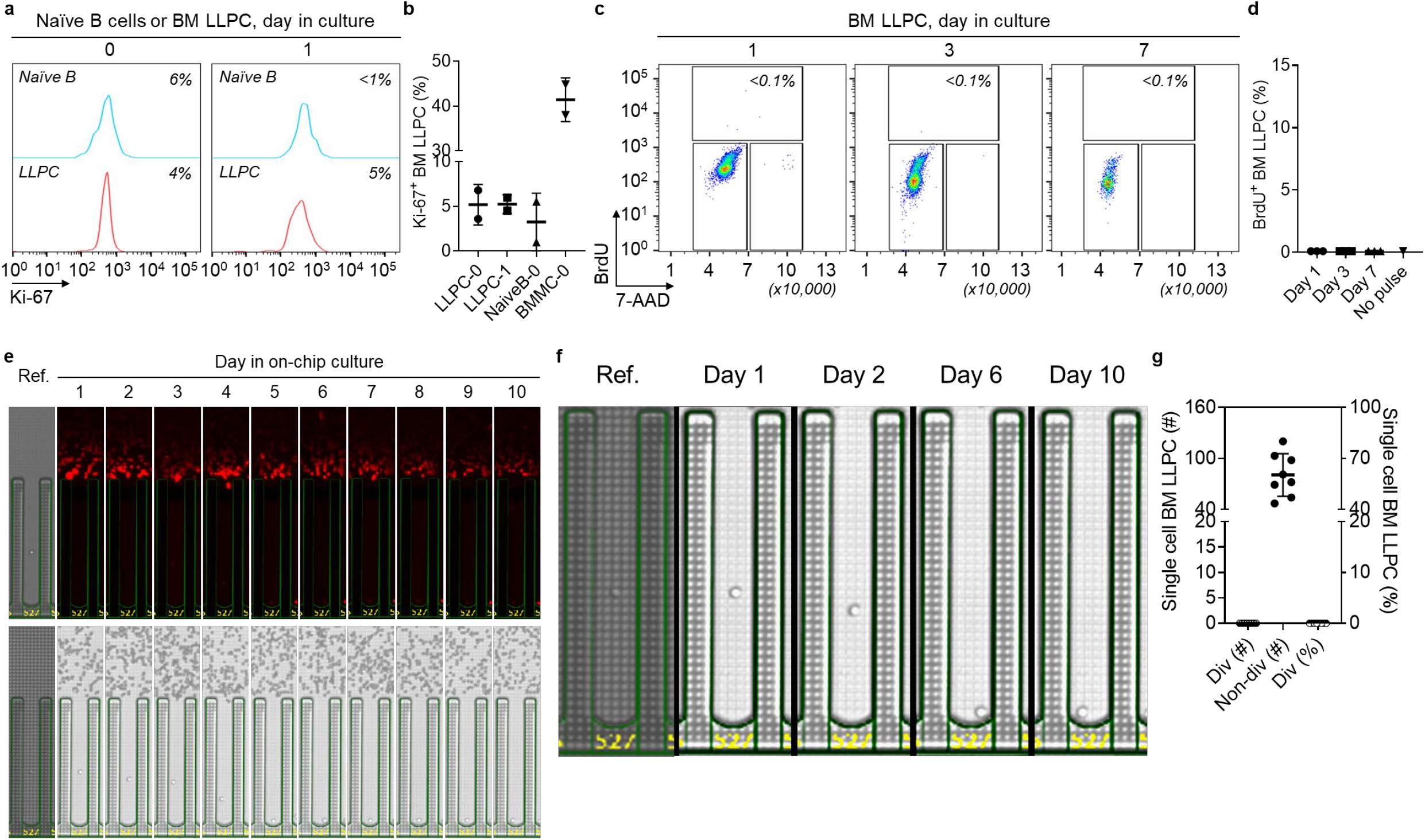
BM LLPC do not divide. (**a**) Representative flow cytometric analysis of Ki-67 expression in BM LLPC at day 0 and 1 in culture (with unstimulated naїve B cells as a non-proliferative control). The numbers indicate the percentages of the cells positive for Ki-67. (**b**) Quantitation of Ki-67^+^ blood ASC (n=2 independent biological replicates). BMMC were used for Ki-67 flow cytometry assay controls (see **Suppl. Fig. S3a**). BMMC, BM mononuclear cells. 0, 1, day in culture. (**c**) Representative flow cytometric analysis of BrdU incorporation in BM LLPC at day 1, 3, and 77 in culture after 3.5-4h pulsing (with apoptotic cells excluded). (**d**) Quantitation of BrdU^+^ BM LLPC pulsed for 3.5-4h (n=3 independent biological replicates). In (**c**): Individual cell cycle phases (S, BrdU^+^; G0/G1, 2N and BrdU^-^; and G2/M, 4N and BrdU^-^) were indicated and the percentage of ASC in S phase was estimated by the amount of BrdU detected. Apoptotic cells were excluded from the analysis. (**e**) Image series displaying a single BM LLPC with ongoing Ig secretion and no evidence of division ((bright field). (**f**) Zoomed in bright-field image series for direct visualization of the non-dividing cell from **(e**) on select days in culture. (**g**) Quantitation (left y axis) and frequency (right y axis) of dividing and non-dividing single cell BM LLPC across 8 independent experiments. Div, dividing; Non-div, no dividing. In (**e,f**): Ref: reference images for visualization of single cell penned at loading.

To assess the proliferation potential of BM LLPC in culture by single cell analysis similar to the blood ASC, BM LLPC were loaded onto the Lightning platform and IgG blooms were captured daily from each single cell over the course of 10-14 days in the PCSS cultures. Not one IgG secreting BM LLPC divided when visualized daily for 2 weeks in culture (**Fig. 2e-g** and **Suppl. Fig. S4a**). Even an immature BM ASC subset, defined as CD19^+^CD38^+^CD138^+^ (PopB)^12,15^ also showed no evidence of proliferation potential in culture by day 4-14 (**Suppl. Fig. S4b,c**). These results confirm human BM CD138^+^ ASC which include the LLPC (PopD) and a less mature BM-resident ASC subset (PopB) do not divide and remain in permanent cell cycle arrest.

## Discussion

In this study, we show that the majority of *bona fide* blood ASC in PCSS culture have stopped dividing despite ongoing expression of Ki-67 and variable BrdU uptake. Using a novel optofluidic single cell analysis platform, we characterized single ASC by secretory function rather than surface markers and found only 11% blood ASC displayed evidence of proliferation. We observed no division after 3 days in culture. While ASC differentiation may be division linked, with upregulation of plasma cell programs especially *PRDM-1* (the gene encoding for BLIMP-1), major hallmarks of cell cycle are downregulated. Although the “blasting” status of an ASC is traditionally used interchangeably with proliferative potential, in the case of early-minted ASC in the periphery, it should be interpreted with caution.

### Upregulation of BLIMP-1 represses c-myc

BCR activation leads to an increase in cell mass and metabolism as activated B cells prepare for proliferation. BCR crosslinking with TLR engagement synergizes to promote clonal expansion and class-switching of antigen-specific B cells^28^. Upregulation of BLIMP-1, a transcription factor that drives B cells to plasma cell differentiation, is known to repress c-myc, a transcription factor that drives functions needed for cell division, thus leading to cell cycle arrest ASC differentiation^29^. However, upstream to ASC differentiation, the DNA breaks involved in class-switching and somatic hypermutation in the germinal center (GC) are critical B cell functions with associated clonal proliferation. As ASC differentiate, the initiation of permanent cell cycle arrest protects from further malignant transformation.

### Role of mTOR in uncoupling differentiation and mitosis

Although ASC differentiation and maturation are ultimately linked to cell cycle arrest, the kinetics of an activated B cell in mitosis and initiation of secretory Ig synthesis is unclear. It was thought that increased Ig production by activated B cells and early ASC requires continuous cell division^30,31^, through which the cells remodel ER and activate the UPR^16^. This mitosis-dependent remodeling is regulated by mTORC1, which controls biochemical pathways generating the cell biomass needed for division^32^. Indeed, during early induction, mTORC1 is crucial for the survival and proliferation of antigen-specific activated B cells in the GC^33^. However, recent reports provide a revised model where activated marginal zone B cells differentiate and adapt to increased Ig production in a cell division-independent process. These marginal zone B cells form ASC with rapid kinetics to generate timely responses to blood-borne pathogens^34^. This alternative model uncouples early ASC differentiation and cell division with mTORC1 proactively “notifying” the UPR^34^. Interestingly, following ASC differentiation, we observe downregulation of mTORC1 pathways as nascent blood ASC mature into LLPC^21^, suggesting possible involvement of mTORC2 signaling.

### ASC cell cycle arrest, cellular senescence, and LLPC maturation programs

In the present study, we confirmed that healthy BM LLPC do not divide for up to 10-14d in culture. BM LLPC have lifelong survival potential but remain “benign” because they engage programs of cell cycle arrest, limiting malignant transformation. Survival without transformation is reminiscent of cellular senescence, which is a stress-induced persistent hypo-replicative state characterized by expression of cell cycle inhibitors^35^. BM LLPC, which enter permanent cell cycle arrest are shown to become resistant to apoptosis^13^ and engage in autophagy pathways^15^. These programs suggest that LLPC may actually undergo cellular senescence programs to prevent malignant conversion as the number of BM LLPC accumulate with age.

### Challenges of assessing proliferation status of human ASC proliferation

Proliferation status of human ASC has been difficult to infer due to three key technical challenges. First, the rapid death of ASC *ex vivo* limits deep investigation into their proliferation profiles. While the time to first division in naїve B cells is typically 24-36h^4,5,36,37^, observation of cell division *in vitro* requires viability for at least 48-72h. The problem studying blood and BM ASC was the rapid apoptosis within hours^21,38-40^. Second, blood and BM ASC are relatively rare. With each infection or vaccination, despite the addition of 10,000-100,000 newly-minted ASC^12,16^, only a small fraction undergoes maturation into LLPC^13^. Of the total BM nucleated cells, the ASC pool represents a tiny fraction (0.1%-0.3%) over a lifetime^41^. Thus, FACS sorting rare ASC populations with total purity has rendered technical limitations for bulk proliferation studies due to contaminating B cells that may undergoing proliferation. Lastly, although FACS sorting is widely used to enrich for ASC, it defines ASC only by surface phenotype and not Ig secretory function, which likely overestimates the number of authentic ASC.

### Cell cycle arrest induced by intrinsic ASC maturation programs versus in vitro culture conditions

Since ASC rapidly die *ex vivo* in conventional cultures, here we used a novel *in vitro* PCSS for both bulk and single cell experiments. The question arises whether the limited proliferation potential of early-minted blood ASC is acquired due to intrinsic characteristics of the cells itself or heavily influenced by the experimentally induced PCSS cultures. We favor the former for several reasons. First, the relationship between *PRDM1* and c-myc and our recent bulk transcriptomics revealed that ASC undergo cell cycle arrest as they mature^13^. Second, our more recent single cell studies with ASC expressing *MKI67* (the gene encoding for Ki-67) cell cycle arrest^42^. In Duan et al.^42^, we show that early BM resident ASC with *MKI67* expression have already upregulated cyclin inhibitors CDKN2A (p16) and CDKN2C (p18) without the use of PCSS. These were found in the same cell in this single cell analysis. Despite *MKI67* expression, it appears that terminal ASC differentiation by its intrinsic nature simultaneously limits further entry into cell cycle.

### Limitation of Ki-67 expression and BrdU incorporation

Assays with Ki-67 and BrdU are often used to infer the proliferative status of mammalian cells. The former is used to assess cells as they exit M phase^43^, while BrdU is used to assess S phase entry and cellular commitment to enter a new cell cycle. In the present study, we used both Ki-67 and BrdU for more comprehensive examination of the ASC proliferation status. However, technically, persistence of Ki-67 proteins after S phase and increased cellular toxicity with longer BrdU labeling led to inconsistent conclusions of the number of proliferating early-minted Ig secreting cells. To our knowledge, the present study is the first to directly visualize both ongoing Ig secretion and cell division of *bona fide* ASC at the single cell level with the use of the novel optofluidic Lightning platform.

### Proliferation of ex vivo ASC versus in vitro ASC generated from B cells

In the absence for antigen, *in vitro* generated effector ASC require cell division to become Ig producing cells^31,44,45^. Earlier work established that the majority (80-85%) of *in vitro* differentiating B cells enters the cell cycle by day 3 and do not undergo cell cycle arrest until day 6^30,31^. Interestingly, these studies also showed that *in vitro* generated ASC must first exit cell cycle for terminal differentiation^30,45^. This study is the first to validate these *in vitro* models by demonstrating that primary *ex vivo* human early-minted ASC or plasmblasts after vaccination also exit cell cycle after terminal differentiation.

In all, our study provides the first single cell visualization of *bona fide* IgG secreting ASC together with direct cell division. Engagement of plasma cell transcriptional programs results in many changes in pathways in cellular structure, processes, and metabolism. However, exit out of cell cycle appears to be essential early step in the maturation process to becoming a LLPC.

## Methods

### Human subjects

Peripheral blood samples were obtained from 64 healthy subjects, 45 of which received vaccines for Tdap, influenza, shingles, meningitis, hepatitis A, hepatitis B, yellow fever, or COVID-19 (the 3^rd^, 4^th^, or 5^th^ dose) at 5-7d prior to sample collection. 17 healthy BM aspirate and 21 femoral head samples were also obtained. Informed consent was obtained from all subjects. All studies were approved by the Emory University Institutional Review Board Committee. All methods were performed in accordance with the relevant guidelines and regulations and in accordance with the Declaration of Helsinki.

### Purification and bulk-culture of blood and BM ASC

PBMC/BMMC isolation was performed as previously described^21^. Fresh blood ASC, BM PopB, and BM LLPC (PopD) were purified using FACS-based sorting, as previously described^15,21^. All sorted blood ASC populations were 88-98% pure. Blood ASC were bulk-cultured in PCSS, which is a culture system that consists of mesenchymal stem cells (MSC) secretome supplemented with APRIL (200ng/mL) and in hypoxic conditions (2.5% O_2_) at 37°C (36°C if for single cell on-chip culture), as previously described^21,22^. BM PopB and BM LLPC (PopD) were bulk-cultured in the same conditions except that no APRIL was supplemented (since exogenous APRIL provides no advantages for survival and secretory functions of human BM ASC in culture)^46^.

### Phase-flow BrdU cell proliferation and flow analysis of Ki-67 intracellular expression

Proliferation of blood ASC and BM LLPC was evaluated on the basis of bromodeoxyuridine (BrdU) incorporation using a phase-flow BrdU commercial kit (BioLegend, Inc.) in accordance with the manufacturer’s recommendations. Briefly, d1, d3, and d7 PCSS-cultured blood ASC were pulsed with 10-15uM BrdU for 3.5-4h, then fixed and permeabilized, and were subsequently treated with DNAse for 1h, followed by incubation with anti-BrdU-FITC antibody (to detect BrdU incorporation) and addition of 7-AAD (10ug/mL; to determine cellular DNA contents; 7-AAD was incubated for 0.5-1.5h prior to acquisition). Negative staining controls for sorted ASC were included to confirm specificity of BrdU (FITC) and 7-AAD staining. FACS-sorted naive B cells were included as non-proliferative controls to establish cell cycle gates. To assess the background binding of the specific anti-BrdU antibody, cultured cells that had not been pulsed with BrdU and were similarly treated were also included. Samples were run on an LSR-II or Symphony A3 flow cytometer and analyzed with the FlowJo v10.8 software (FlowJo, LLC). The percentage of ASC in S phase was estimated by the amount of BrdU-FITC detected.

For intracellular staining of Ki-67, cells were fixed and permeabilized, followed by incubated with anti-Ki-67 antibody or isotype controls. Cellular fluorescence data was acquired with a Symphony A3 flow cytometer and analyzed with the FlowJo v10.8 software (FlowJo, LLC).

### Single cell Ig secretion and proliferation analysis

#### Lightning optofluidic system preparation

For functional single cell analysis, the Lightning system (Berkeley Lights, Inc.; now PhenomeX, Inc.), equipped with the OEP technology which can identify single ASC and precisely position them individually into nanoliter-volume pens (i.e. a NanoPen chamber), was used. The system was sterilized by purging all fluidic lines before the initiation of the workflows.

#### Blood and BM ASC single cell imports

After sterilization, an OptoSelect 1500 chip was loaded, wetted, primed, and focus-calibrated using standard workflows. Immediately after FACS sorting, a volume of 10uL cell suspension containing 3,000-25,000 FACS-purified ASC in PCSS was imported into the chip channels through a customized Targeted Pen Selection (TPS) penning algorithm with a small volume import (SVI) operation – and with or without penning chip by multiple rounds of imports and/or by fields of view (FOV) indexing. Following equilibration, the system performed reference imaging by capturing the bright-field high-magnification (4x and 10x) images across the chips (as the post-penning background reference and the normalization reference for ASC quantity during on-chip culture). Single cell ASC were penned in fresh PCSS supplemented with Loading Reagent (Berkeley Lights, Inc.; now PhenomeX, Inc.). Residual cells in the channels were subsequently flushed to a sterile PCSS collection tube for reuse.

#### On-chip blood and BM ASC single cell cultures

Following single cell deposition into NanoPens, ASC were cultured on-chip at 36°C with constant perfusion of fresh PCSS at a rate of 0.01μL/s and in hypoxic condition that was infused with a pre-analyzed gas mixture containing 2.5% O_2_, 5% CO_2_, and 92.5% N_2_ (AirGas). To keep track of penned ASC numbers, timelapse bright-field images were taken daily to keep track of cell growth for each colony. For blood ASC, PCSS was supplemented with APRIL (200ng/mL); for BM ASC, all but one were without APRIL.

#### Bead-based, in-channel IgG capture assay

In-channel IgG capture assays were performed using single assay mixtures containing anti-human IgG-coated 6-8μm beads (Spherotech) and fluorescently-labeled detection anti-IgG antibodies (2.5μg/mL; Jackson ImmunoResearch). The mixtures were imported into the channels and floated above the pens throughout the course of the capture assay. IgG secreted by single cells contained in pens diffused into the channels and bound the beads. Accumulation of fluorescence from secondary antibodies on the IgG-coated bead upon secreted IgG binding led to the development of fluorescent halos – or blooms – in the channels adjacent to the pens. In this way, secreted IgG concentrations (which represent ASC secretion rate) could be semi-quantitatively assessed. IgG bloom development was time-dependent, with capture assays running for 60-80 minutes, and imaged at 10-minute intervals using a TRED (AF-594), FITC (AF-488), or PE (AF-555) filter cube. Single, penned ASC were directly captured for functional Ig secretion (as “blooms”) and simultaneously enumerated for the quantity of cells by direct bright field microscopy. The chip was flushed extensively after each assay (to remove all bead assay mixture). All assays were performed with PCSS.

#### Workflows and chip image analysis

All the workflow operations were controlled using the Cell Analysis Suite (CAS) software (v2.2.8.40; Berkeley Lights, Inc.). IgG blooms and ASC quantitation were determined using Image Analyzer (v2.2; Berkeley Lights, Inc.) and Assay Analyzer (v2.2; Berkeley Lights, Inc.), and subsequently confirmed through manual assessment.

## Acknowledgements

We thank Drs. Jonathan Didier, Suhagi Baxi, and Eugene Gibbs of Berkeley Lights, Inc. (now PhenomeX, Inc.), and Shuya Kyu and Ariel Ley for technical assistance. We thank Robert E. Karaffa, Kametha T. Fife, and Sommer Durham of the Emory University School of Medicine Flow Cytometry Core (EFCC) for technical support. We also thank our team of clinical coordinators and donors who made this study possible.

---

**Fig. S1.**
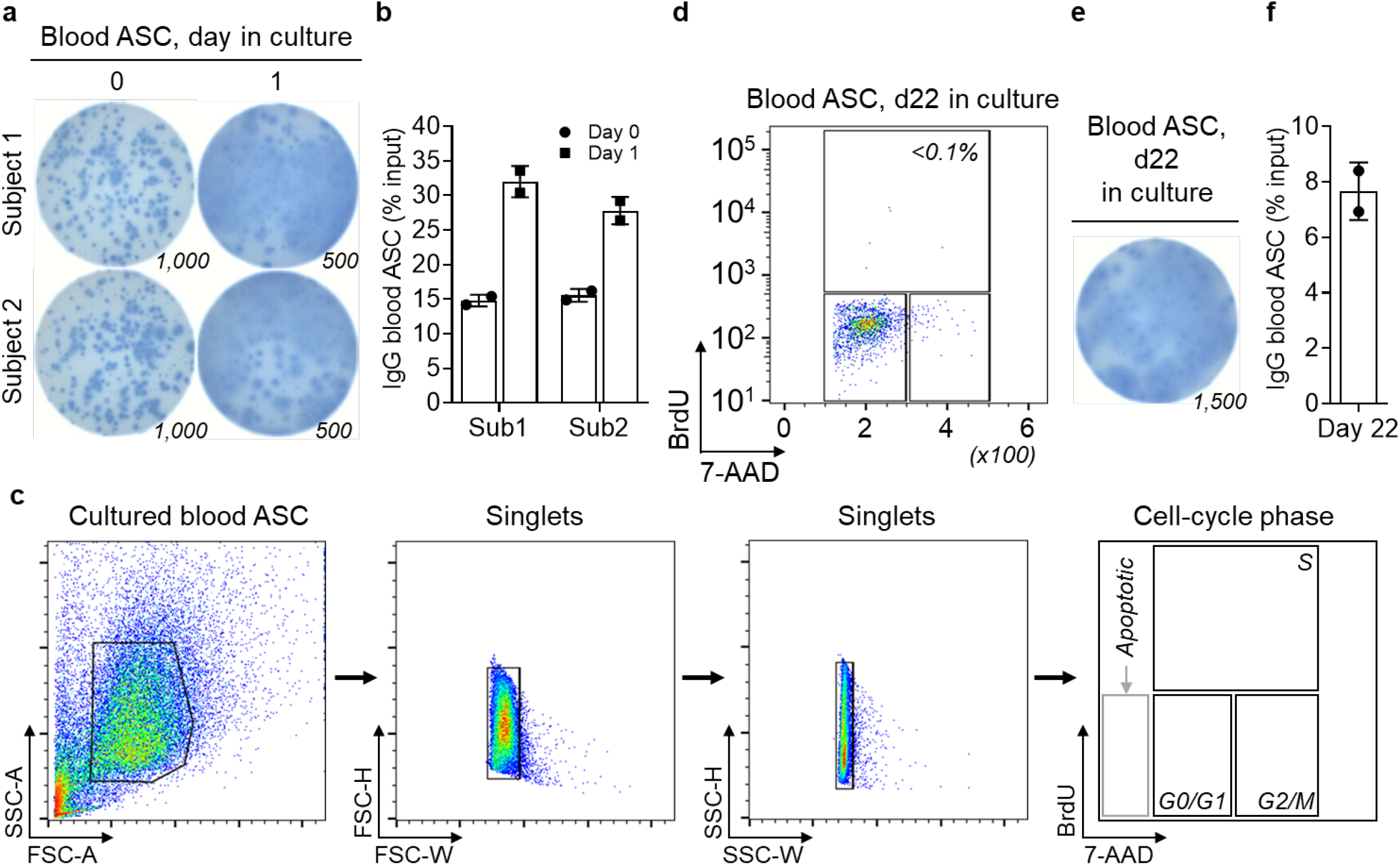
BrdU flow cytometric analysis of cultured blood ASC and bulk IgG ELISpots. (**a**) Cultured blood ASC were assayed for IgG secretion by ELISpots. (**b**) Quantitation of IgG secretory ASC in (**a**). (**c**) General gating strategy used for flow cytometric analysis to assess cell cycle phase in cultured blood ASC. The end panel represents a diagram for phase-flow BrdU cell proliferation with the individual cell-cycle phases (S, BrdU+; G0/G1, 2N and BrdU-; and G2/M, 4N and BrdU-) indicated. (**d**) Flow cytometric analysis of BrdU incorporation in blood ASC at day 22 in culture. The percentage of ASC in S-phase was estimated by the amount of BrdU detected (analyzed with apoptotic cells excluded). (**e**) Blood ASC at day 22 in culture were assayed for IgG secretion by ELISpots. (**f**) Quantitation of IgG secretory ASC in (**e**). In (**a,e**): Representative IgG ELISpot images shown. The number below indicates the calculated input number of ASC that were measured at day 0.

**Fig. S2.**
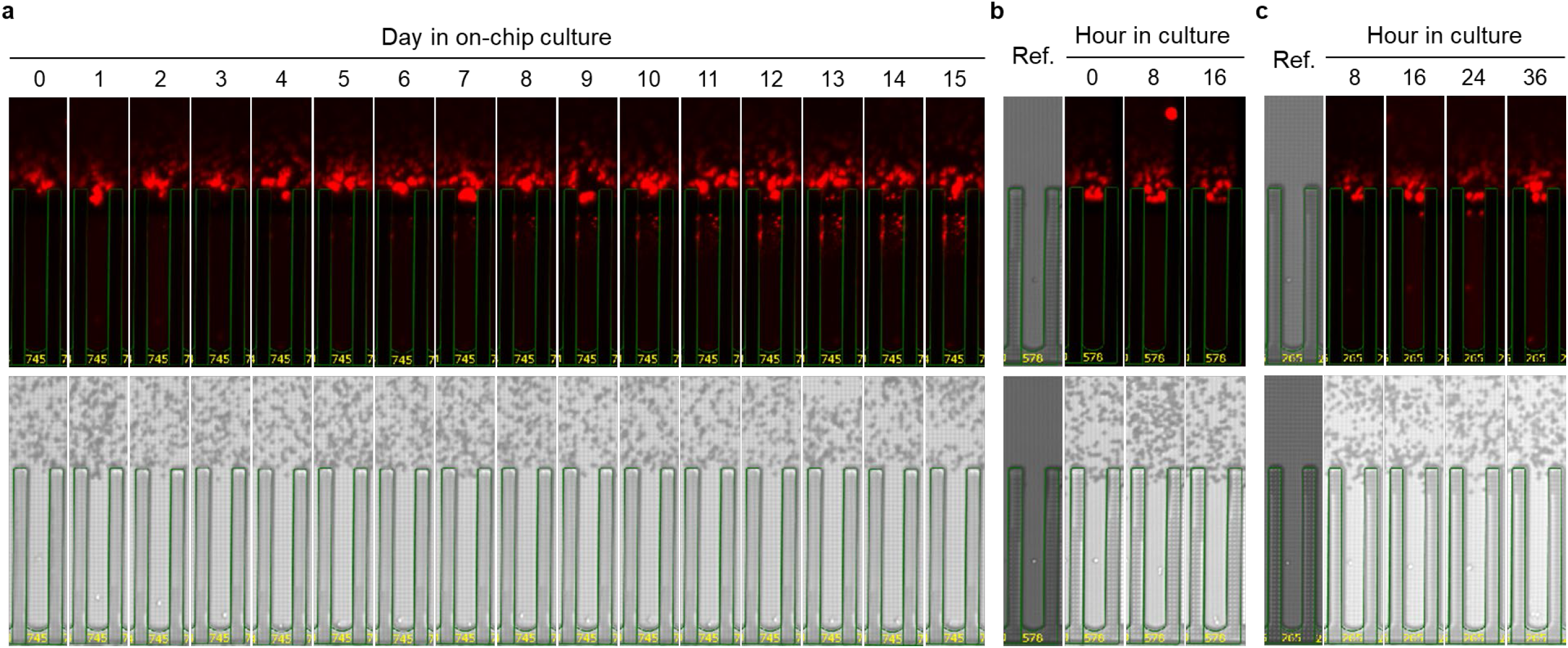
Timing of the few IgG secreting blood ASC that divide. (**a**) Majority of blood ASC do not divide when observed to day 15. Image series displaying a representative single blood ASC with ongoing Ig secretion and no evidence of division ((bright field) as far as day 15. (**b,c**) Representative blood ASC undergoing division within 8h (**b**) or 36h (**c**). Cell division was assessed by visualization of cell addition ((bright field) over the course of the culture. In (**b,c**): Ref: reference image show single cell penned at time of loading.

**Fig. S3.**
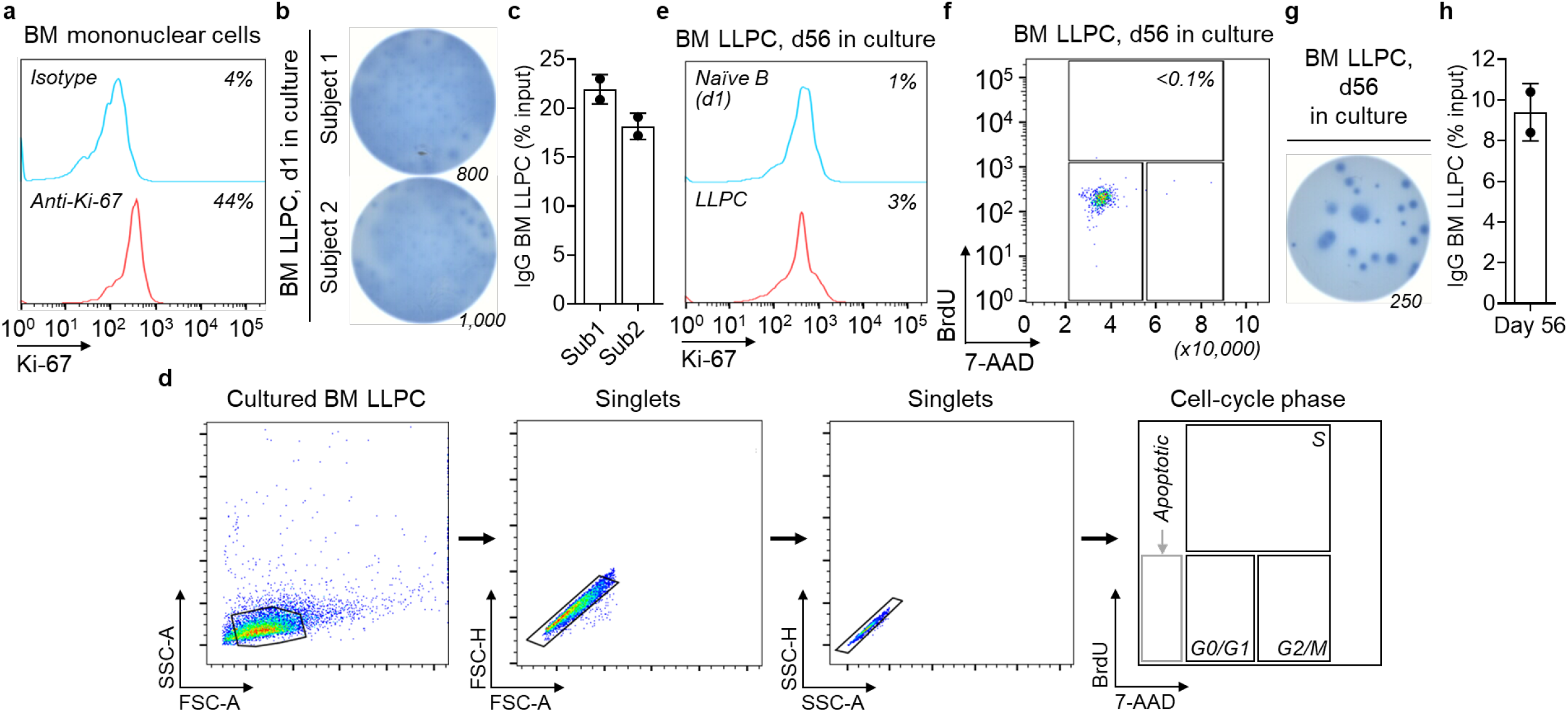
BrdU flow cytometric analysis of cultured BM LLPC and bulk IgG ELISpot assays. (**a**) Ki-67 flow cytometry assay controls using BMMC were stained with isotype controls or anti-Ki-67 antibody. The numbers indicate the percentages of the cells positive for Ki-67. (**b,c**) IgG ELISpots of cultured BM LLPC. Quantitation of IgG secretory BM LLPC in (**b**). (**d**) Gating strategy used for flow cytometric analysis to assess cell cycle phases in cultured BM LLPC. The last panel represents a diagram for phase-flow BrdU cell proliferation with the individual cell-cycle phases (S, BrdU+; G0/G1, 2N and BrdU-; and G2/M, 4N and BrdU-) indicated. (**e**) Flow cytometric analysis of Ki-67 expression in BM LLPC at d56 in culture (with unstimulated naїve B cells at d1 as a non-proliferative control). The numbers indicate the percentages of the cells positive for Ki-67. (**f**) Flow cytometric analysis of BrdU incorporation in BM LLPC at d56. The percentage of ASC in S-phase was estimated by the amount of BrdU detected (analyzed with apoptotic cells excluded). (**g, h**) BM LLPC at d56 were assayed for IgG secretion by ELISpots. Quantitation of IgG secretory BM LLPC in (**g**). In (**b,g**): Representative IgG ELISpot images shown. The number below indicates the calculated input number of ASC at day 0.

**Fig. S4.**
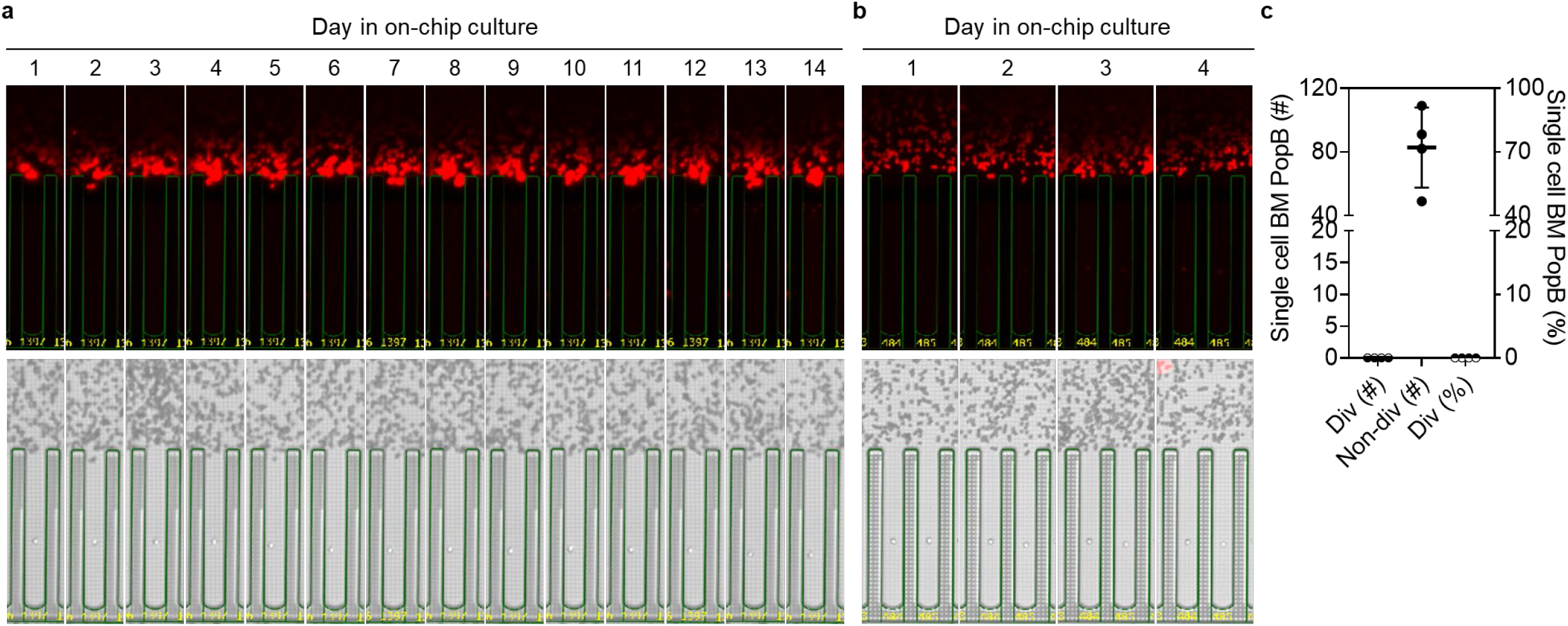
Mature BM ASC with Ig secretion do not divide even up to day 14 in single cell cultures. (**a,b**) Image series displaying a single BM PopD (LLPC) (**a**) or BM PopB (**b)**; 2 nearby pens each with a single ASC) with ongoing Ig secretion showing no evidence of division ((bright field) by day 4. (**c**) Quantitation (left y axis) and frequency (right y axis) of dividing and non-dividing single cell BM PopB across 4 independent experiments. Div, dividing; Non-div, no dividing. In (**a,b**): Cell division was assessed by visualization of cell addition ((bright field) over the course of the culture. In (**c**): Div, dividing; Non-div, no dividing.

